# Improving the power of drug toxicity measurements by quantitative nuclei imaging

**DOI:** 10.1101/2023.12.28.573573

**Authors:** Alesya M. Mikheeva, Mikhail A. Bogomolov, Mikhail V. Sementsov, Pavel V. Spirin, Vladimir S. Prassolov, Timofey D. Lebedev

## Abstract

Imaging-based anticancer drug screens are becoming more prevalent due to development of automated fluorescent microscopes and imaging stations, as well as rapid advancements in imaging processing software. Automated cell imaging provides many benefits such as their ability to provide high-content data, modularity, dynamics recording and the fact that imaging is the most direct way to access cell viability and cell proliferation. However, currently most publicly available large-scale anticancer drugs screens, such as GDSC, CTRP and NCI-60, provide cell viability data measured by assays based on colorimetric or luminometric measurements of NADH or ATP levels. Although such datasets provide valuable data, it is unclear how well drug toxicity measurements can be integrated with imaging data. Here we explored the relations between drug toxicity data obtained by XTT assay, two quantitative nuclei imaging methods and trypan blue dye exclusion assay using a set of four cancer cell lines with different morphologies and 30 drugs with different mechanisms of action. We show that imaging-based approaches provide high accuracy and the differences between results obtained by different methods highly depend on drug mechanism of action. Selecting AUC metrics over IC50 or comparing data where significantly drugs reduced cell numbers noticeably improves consistency between methods. Using automated cell segmentation analysis and data-driven approach we show that XTT assay produces unreliable data for cell cycle and DNA repair inhibitors due induced cell size growth and increase in mitochondria activity. We also explored several benefits of image-based analysis such as ability to monitor cell number dynamics, dissect changes in mitochondria activity from cell proliferation, and ability to identify chromatin remodeling drugs.

## Introduction

Large-scale drug screens provide valuable data for understanding drug mechanisms of action^1,2^, cancer cell vulnerabilities^3^, development of novel drugs^4,5^ and drug repurposing^6^. The ability to kill specific cancer cells is a conventional indicator of anticancer drugs efficacy, however studies often rely on different methods to measure cell viability. Thus, several largest datasets provide drug toxicity data measured by methods, which rely on NADH activity, ATP or protein levels: Genomics of Drug Sensitivity in Cancer (GDSC) uses resazurin and CellTiter-Glo^3^, Cancer Therapeutics Response Portal uses CellTiter-Glo^1^, and NCI-60 uses sulforhodamine B assay^7^. These methods rely on indirect measurement of drug toxicity, and sometimes can be unreliable because drugs may influence cellular metabolic activity or protein levels without change in cells quantity^8,9^. Direct counting-based methods, such as trypan blue dye exclusion, are used to identify number of surviving cells. However, such methods constitute a laborious task for an operator, may give non-reproducible results and are not suitable for large-scale studies. Novel drug screens utilize more direct approaches such as measurement of DNA-barcoded cells in PRSIM study^10^ or real-time measurements of cell occupied area by IncuCyte or xCELLigence^11,12^. Data obtained by different methods may have poor agreement^13,14^ due to changes in cell metabolism^8^, adhesion, cell size and morphology^11^, or drug influence on substrate used for measurement. Optimization the consistency between NADH or ATP-based assays, measurements of cell area or LIVE/DEAD assays was addressed by numerous studies, providing either protocol optimizations, metric selection or method combination^15-18^.

Advances in microscopy and image processing algorithms led to new high-content screening methods using fluorescence microscopy^19^, that also allow viable cells counting^20-25^. Microscopy provides the most direct approach to measure cell proliferation and drug toxicity^20^, as it does not require extensive processing of cells, such as trypsinization and cell lysis, highly customizable by the use of fluorescent stains and proteins which allows measurement of cell cycle and cell death^26^, protein activity^27-29^, cell differentiation^30^, metabolites levels and cellular morphology^31^. However, accurately segmenting single cells or nuclei is still a difficult task due to different growth patterns and drug-induced morphological changes.

Although relations between most common cell viability measurement methods were vastly explored before, there is still little data on how cell viability measurements using imaging-based methods correspond with other cell viability assays. Such data is essential for reliable integration of drug screen data from studies that used different readout methods and for selecting the appropriate study design. Thus, the main aim of this study was to compare cell viability measurements made by XTT colorimetric assay, which is similar to MTT and WST, but does not require solubilization step, trypan blue dye exclusion and quantitative imaging to provide proper solutions for integrating data obtained by different methods. Do determine number of cells by fluorescence microscopy we stained cell nuclei with Hoechst-33342 or utilized cells with continuous expression of fluorescent H2B-mRuby protein that allows automated nuclei counting. Since drug mechanisms of action and cell morphology can influence imaging results^32^ we utilized different anticancer drugs and cancer cell lines. Here we explore how the mechanisms of action for different drugs affect differences in viability measurements performed by different methods and whether the results depend on a cell type used.

## Results

### XTT measurements significantly differ from quantitative nuclei imaging

To determine differences between some assays for measuring cell viability cancer cell lines of various origins: lung adenocarcinoma H1299, glioblastoma LN-18, ovarian adenocarcinoma SK-OV-3 and neuroblastoma SH-SY5Y were treated by different inhibitors and measured cell viability using four different methods (Figure 1A). We used previously established H1299 and SH-SY5Y cells with H2B-mRuby expression^29^ and introduced H2B-mRuby marker to LN-18 and SK-OV-3 cells by lentiviral transduction. Since not all cell population can be uniformly transduced (Figure 1B), which may affect assay results, we also used imaged-based counting using nuclei staining with Hoechst-33342 (Figure 1A). Trypan blue dye exclusion assay (TB) allowed us to determine the exact quantity of viable cells and was later used as a benchmark. In order to account for different cell death mechanisms induced by anticancer drugs we selected six commonly used drugs with various mechanisms of action: doxorubicin, etoposide, dasatinib, gefitinib, panobinostat and azacitidine (5-Aza).

**Figure 1.**
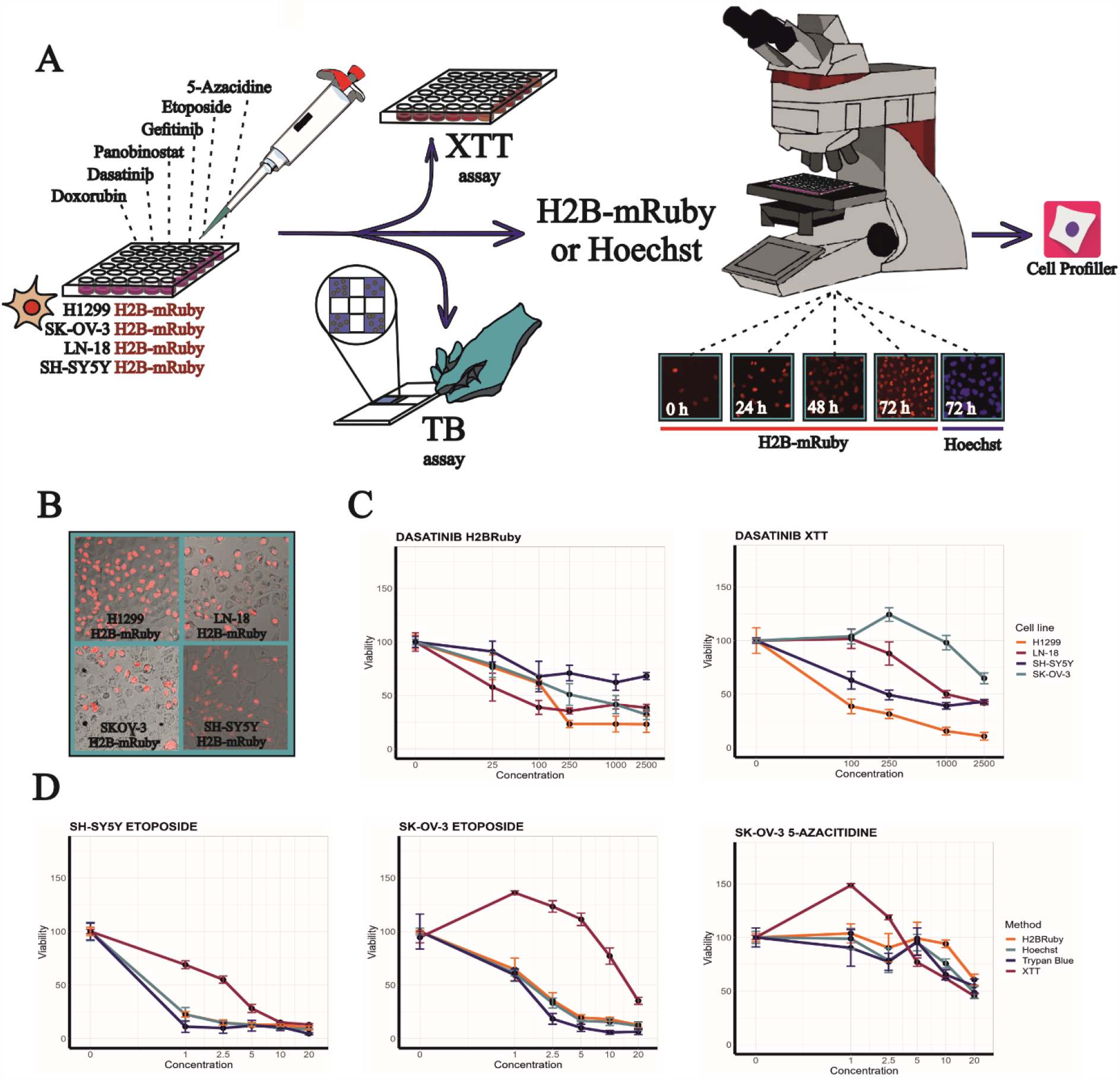
Cell viability measurement by different methods. **A**. Experiment design scheme. TB-trypan blue exclusion assay. **B**. Representative images of cells, expressing H2B-mRuby protein (red). **C-D**. Dose-dependent changes in cell viability measured at 72h after drug treatment by XTT, trypan blue exclusion and nuclei counting using Hoechst staining or H2B-mRuby cells. Data presented as means and SD values for four repeats. Data was grouped by different cell lines for a single method **(C)** or by different methods for a single cell line **(D)**. Concentrations are provided in logarithmic scale. Dasatinib was used in 25-5000 nM range, etoposide and 5-azacytidine in 0.25-20 μM range. All data was normalized on cell viability measurements for cells mock-treated with 0.1% DMSO.

The difference between measurements obtained by different methods varied depending on a cell line and drug used. For example, the difference between SK-OV-3 and H1299 sensitivity to dasatinib was much higher when measured by XTT than using H2B-mRuby nuclei counting (Figure 1C). The XTT assay produced exaggerated cell viability values in contrast to the results obtained by other methods, as observed for all cell types treated with etoposide or 5-Aza (Figure 1D). However, in the case of 5-Aza, the difference in response was observed primary by non-toxic drug concentrations were XTT measurement provided higher readouts than in DMSO control (zero drug concentration).

### Variance in IC50 values is caused by combination of particular drug and method

To quantitatively compare cell viability data we calculated IC50 (as concentration at which cell viability is reduced to 50%) and AUC values and then examined correlations between values obtained by different methods. Correlation between IC50 values obtained from TB assay and H2B-mRuby or Hoechst nuclei counting were significant (r = 0.76 and 0.72; p-values < 0.0001) (Figure 2A). Correlations between XTT and TB or H2B-mRuby assays were not significant (r = 0.28 and 0.36; p-values > 0.08). However, when we compared the AUC values, correlations between each method pair were significant (Figure 2B), even for TB and XTT assays (r = 0.76; p-value<0.0001), which had the weakest correlation for IC50 values (Figure 2A). In general, comparing AUC values dramatically improved correspondence between data obtained by different methods (Figure 2B). To investigate whether the cell origin or drug type drives the differences between IC50 values for selected methods, we compared differences in IC50 values for each treatment with mean IC50 values for a particular drug (Figure 2C). As the result, we observed that XTT results had the highest number of identified outliers (at least two-fold change in IC50) (Figure 2 C-D). Seven out of eleven outliners among all drugs and cell lines were for dasatinib, suggesting that a particular method-drug combinations has higher influence on differences between measurements than the cells origin. We had not detected any outliners for AUC values, meaning that AUC values have less dependency on a method used to determine cell viability.

**Figure 2.**
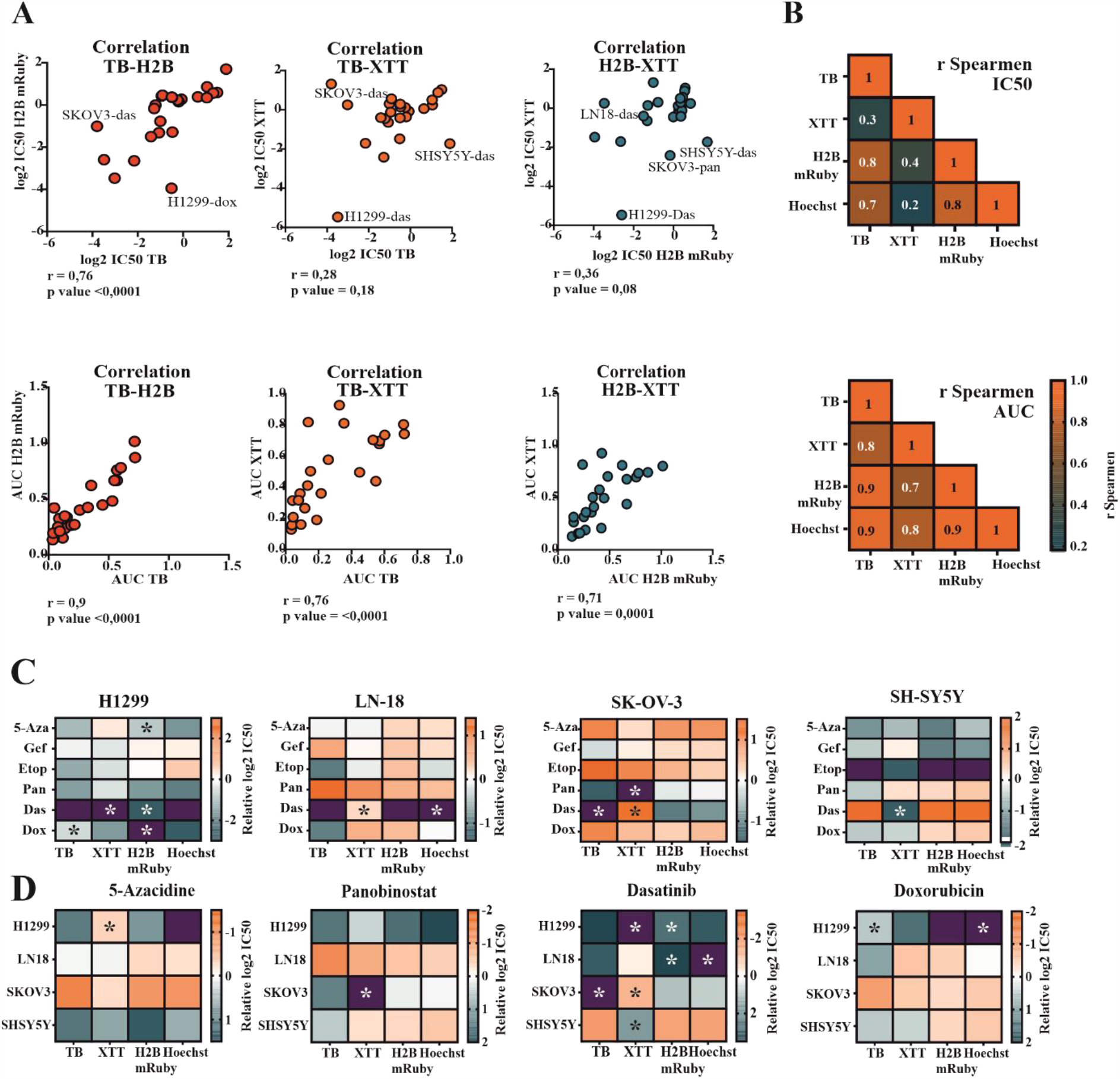
Correlation of IC50 and AUC between different assays. **A**. Dot plots comparing IC50 and AUC values for different methods. Each point corresponds to cells treated with particular drug, most prominent outliners are annotated. Spearman’s correlation R and p-values are provided under each graph. H2B-values obtained by nuclei counting using H2B-mRuby, TB-trypan blue exclusion assay, das-dasatinib, dox-doxorubicin, pan-panobinostat. **B**. Heatmap of pairwise Spearman’s correlations for IC50 and AUC values obtained by different methods. Insignificant correlations are highlighted by blue color (p<0.05). **C-D**. Log2 IC50 differences heatmaps. For each cell lines and drug respective IC50 values were normalized on mean IC50 value for that drug across all cell lines. IC50 values were grouped by cell lines **(C)** or by drug **(D)**. Color shows log2 difference between IC50 value and the mean IC50 for that drug and thus shows how sensitive particular cell line to a drug. Differences higher than 4-fold are marked as purple. Stars highlight cases when the differences are more than two-fold.

Since drugs had significant effect on measurement variation between methods, we additionally compared measurements obtained by H2B-mRuby imaging and an XTT assay using H1299 cells and a panel of 30 drugs with different mechanisms of action. Overall, the Spearman’s correlation between methods was significant (r=0.77; p-value < 0.0001) (Figure 3A). The highest differences between methods were caused by cell cycle inhibitors palbociclib, alisertib and adavosertib, DNA-damage repair inhibitor talazoparib and PKC inhibitor staurosporine. We observed the most difference between measurements were caused by lower drug concentrations that reduced cell proliferation less than by 50%. For some drugs, for example palbociclib or talazoparib, XTT assay failed to detect increasing drug toxicity (Figure 3B).

**Figure 3.**
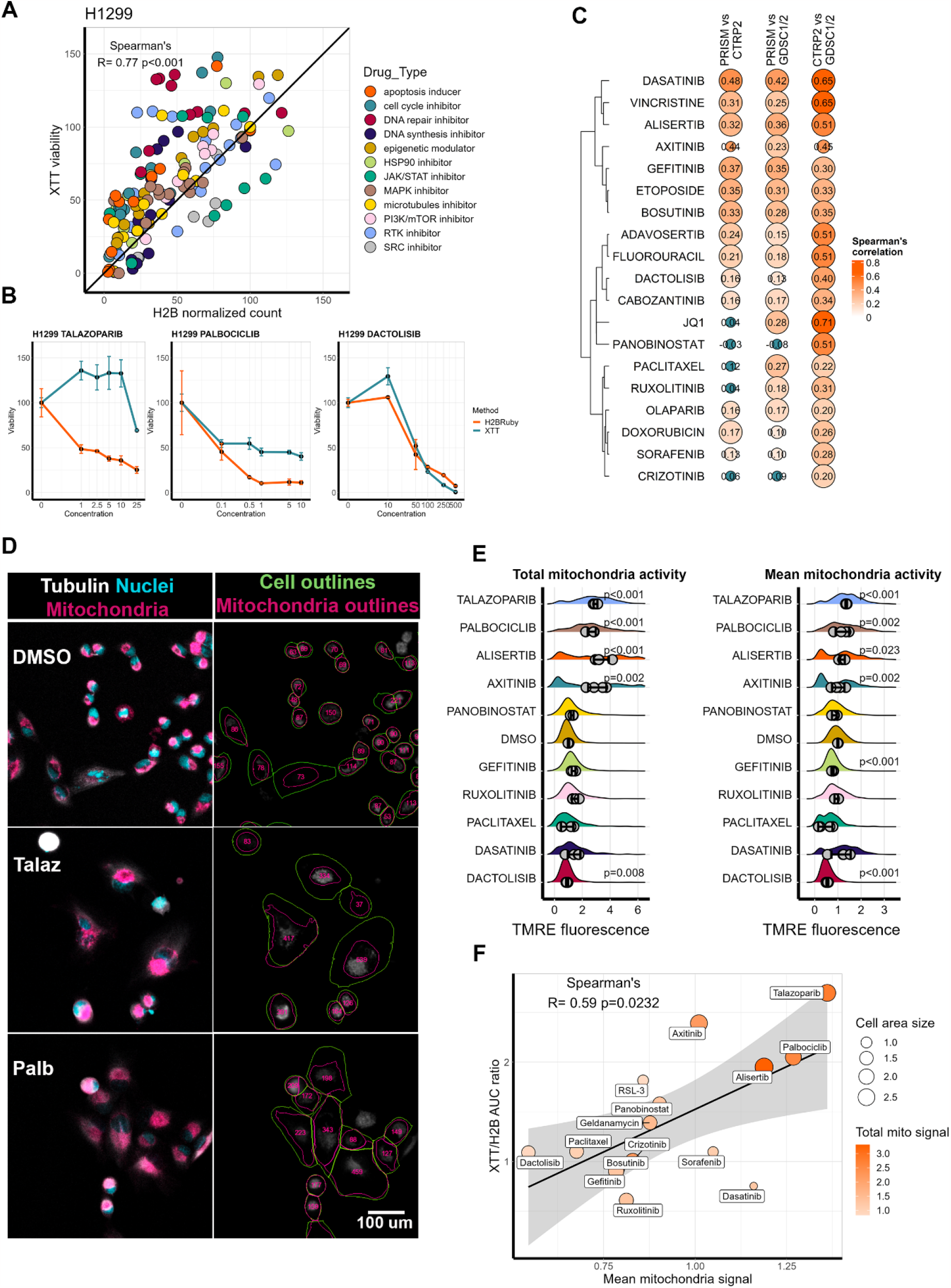
Drugs affect XTT measurements depending on their mechanism of action. **A**. Correspondence between normalized cell viability of H1299 cells measured by XTT and H2B-mRuby imaging. Each dot represents the mean values between three repeats for each drug concentration. Cell viability values were normalized on values for cells mock-treated with DMSO. **B**. Dose-dependent response of H1299 to talazoparib (1-25 μM), palbociclbib (0.1-10 μM) and dactolisib (10-500 nM). **C**. Spearman’s correlations of AUC values between PRISM GDSC1/2 and CTRP2 datasets. **D**. Staining and segmentation of H1299 cells treated with DMSO, palbociclib (Palb) or talazoparib (Talaz) for 72h. In live cells tubulin was stained by Tubulin Tracker DeepRed (gray), nuclei with Hoechst-33342 (blue), and mitochondria with TMRE (magenta). Cell borders and areas occupied by mitochondria in each cell were determined by Cellpose and CellProfiler software. Total mitochondria signals (integrated TMRE fluorescence) for each cell are indicated by numbers. **E**. Distribution of integrated TMRE signal per cell (total mitochondria activity) and TMRE signal per cell normalized to area occupied by mitochondria (mean mitochondria activity) in H1299 cells treated with drugs or DMSO for 72h. For each drug data for two toxic drug concentrations was combined. Dots show mean values for each biological repeat (n=4) and SD based on repeats is provided. For each repeat four automatically selected imaging fields were analyzed. On average 650 cells were used to calculate each distribution. P-values were calculated using Mann-Whitney test by comparing mean values for each image with DMSO. **F**. Correlation between the ratio of AUC obtained from XTT assay measurements to H2B-mRuby imaging and drug effect on mean mitochondria activity normalized to area occupied by mitochondria. Size of each dot is proportional to mean cell area as calculated by CellProfiler and color is proportional to total mitochondria activity induced by a drug.

To verify that similar relationship between methods is relevant for high-throughput drug screens and not unique to our case we compared drug sensitivity data for the drugs from three databases: GDSC1/2^3^, CTRP^1^ and PRISM^6^. Cell viability in GDSC1/2 and CTRP were measured by colorimetric or luminometric assays: resazurin and Syto60 in GDSC1, and CellTiter-Glo in GDSC2 and CTRP. In PRISM dataset cell lines were labeled by DNA-barcodes and then pooled drug assays were performed and cell proliferation was measured by sequencing and enumerating the number of DNA-barcodes. As expected GDSC1/2 and CTRP datasets had high correlation scores, however PRISM had considerably weaker correlations with both GDCS1/2 and CTRP. Only 7 drugs had R > 0.3 when PRISM data was compared with GDSC1/2 and 5 drugs did not have significant correlation for one of the comparisons (Figure 3C). Notably the weakest correlations were for panobinostat and ruxolitinib, which also demonstrated high difference between measurement methods in our study.

### Difference in cell viability measurements depends on drug-induced mitochondrial activity

As other studies suggested the elevated results of cell viability assays can be caused by increased mitochondria activity or by drug directly affecting substrate. Thus, we selected 15 drugs with different effects on XTT and stained H1299 cells 72h after drug treatment tubulin and nuclei to determined cell morphology and for mitochondrial activity using potential-dependent TMRE fluorescent stain. Each drug was used in two concentrations which inhibited cell proliferation and single cells were automatically segmented using custom pipeline. Several drugs, such as palbociclib, talazoparib, alisertib and axitinib significantly increased cell size and overall TMRE staining intensity (Figure 3D, E). Moreover, we detected an increase in TMRE signal when it was normalized by area occupied by mitochondria (Figure 3E), meaning that these drugs not only increase overall cell size and thus mitochondria number, but also mitochondrial increase activity. We did not detect such increase in mitochondrial activity for cell treated with other drug at toxic concentrations, except for dasatinib, meaning that this effect is specific to certain drugs. TMRE fluorescence normalized by area positively correlated (r = 0.59; p-value = 0.02) with ratios between AUC values obtained by XTT assay and H2B-mRuby imaging (Figure 3F). We also tested the effect of drugs added in growth medium without cells on absorption in XTT assay due to drug absorption properties or interaction with XTT, but we did not detect any significant changes in the readouts. These data provide systematic verification that selective increase in mitochondria activity caused by certain drugs affects readouts by metabolic-based assays.

### H2B-mRuby imaging reveals different cell death dynamics

The use of H2B-mRuby provides a non-invasive way to observe the dynamics of cell proliferation throughout the experiment. Although other methods like XTT or Hoechst staining also allow to measure dynamics they can significantly affect cellular processes and may cause additional toxicity. The dynamic analysis of cell viability revealed several types of responses based which couldn’t be predicted by endpoint analysis at 72h. Our results highlight that interpretation of IC50 values at defined time point should take into account proliferation rate of particular cell line. For example, although 5-Aza at 5 μM reduced SH-SY5Y cell numbers by two times, it failed to prevent cell proliferation (Figure 4A) and actual proliferation inhibition occurred only at 20 μM. On the other hand, for SK-OV-3, which have slower proliferation rate, the two-fold drop in cell number at 72h indicates full proliferation inhibition (Figure 4B). In some cases, measuring cell number dynamics can help to distinguish drugs that actively kill cells and not just slow down proliferation. For example, we detected reduction in number of nuclei for SH-SY5Y treated with etoposide from 24h and for SK-OV-3 treated with gefitinib or doxorubicin after 48h. All together our observations suggest that monitoring cell proliferation dynamics may be useful to better discriminate drugs with different mechanisms of action (Figure 4C, D). However, the use of fluorescent protein may be restricted by drug fluorescence, such as in case of 2500 nM of doxorubicin, which caused a false increase in cell numbers at 24 and 48h due to high accumulation of fluorescent drug in cells (Figure 4B).

**Figure 4.**
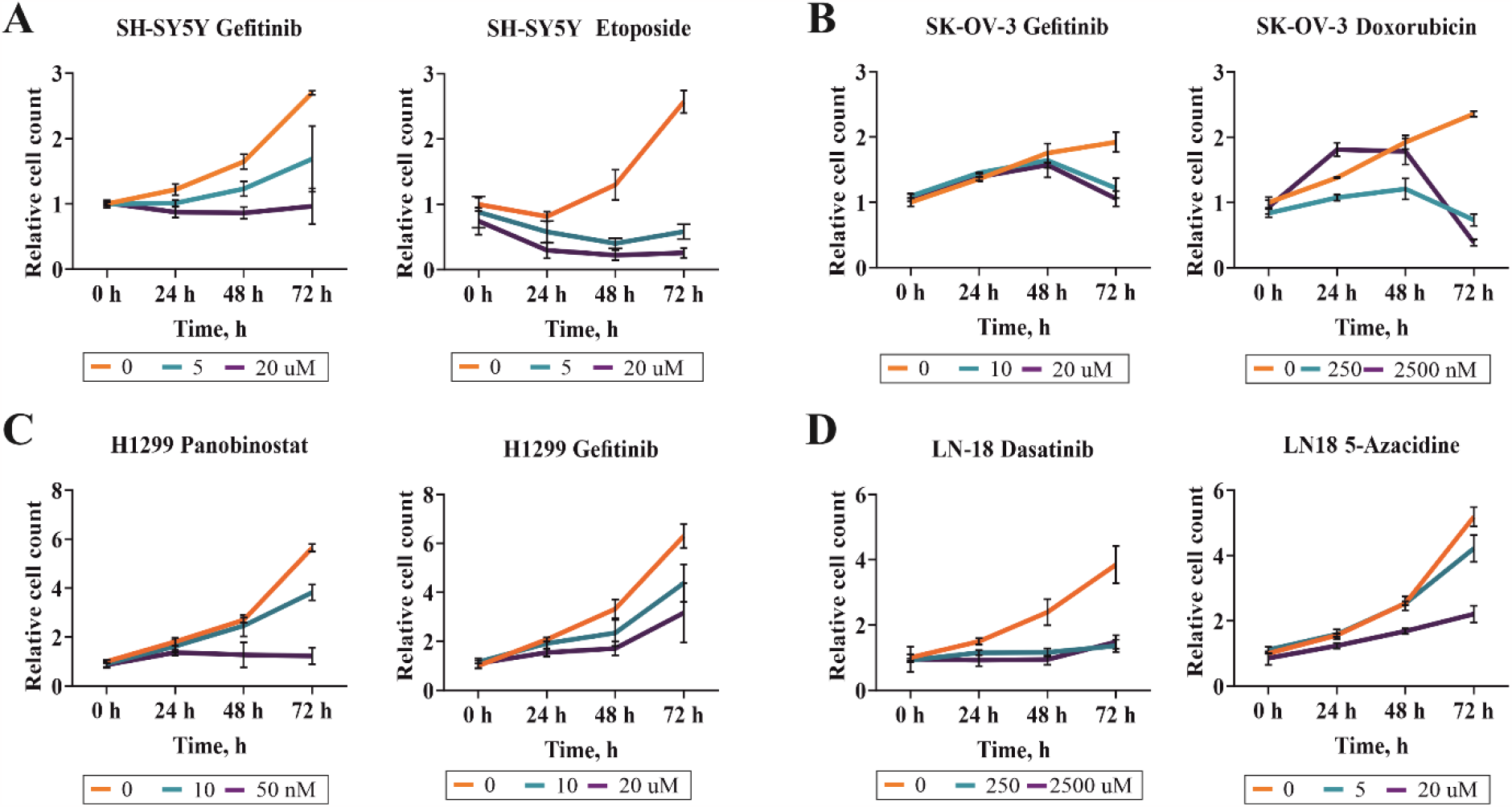
Measurement of cell proliferation dynamics using H2B-mRuby. Graphs show change in a number of identified nuclei for different drug concentration depending on time after start of the treatment. Data presented as means and SD values based on four repeats. Nuclei counts were normalized to the nuclei counted before the start of the treatment.

### H2B-mRuby fluorescence can be used to identify chromatin remodeling drugs

When we processed H2B-mRuby images, we also noted that several drugs caused an increase in H2B-mRuby fluorescence intensity. Across all cell lines, this increase was prominent when cells were treated with panobinostat and this effect was concentration-dependent (Figure 5A, B). We hypnotized that increase in H2B-mRuby fluorescence may be due to increase in H2B-mRuby expression after HDAC inhibition by panobinostat. Viral promoters often get silenced in cells and HDAC inhibitors are known to be able to reactivate HIV-1 gene expression during latent infection stage^33^. To verify, that this effect was caused by HDAC inhibition we additionally tested another HDAC inhibitor belinostat. It increased H2B-mRuby signal intensity in a similar manner as panobinostat (Figure 5B). To check if this effect is specific to HDAC inhibitors we measured H2B-mRuby intensity under all 30 tested drugs for H1299 cells. We found that well described chromatin remodeling drug JQ-1 that inhibits BET also significantly increased H2B-mRuby fluorescence (Figure 5B). All other drugs, except HSP90 inhibitors geldanamycin and 17-DMAG, did not have significant effect on H2B-mRuby signal intensity (Figure 5C). None of the drugs expect doxorubicin were fluorescent by themselves in used concentrations as was tested on H1299 cells without H2B-mRuby. These findings suggest that a fluorescent protein expressed as a transgene can be additionally used to find drugs with chromatin remodeling properties.

**Figure 5.**
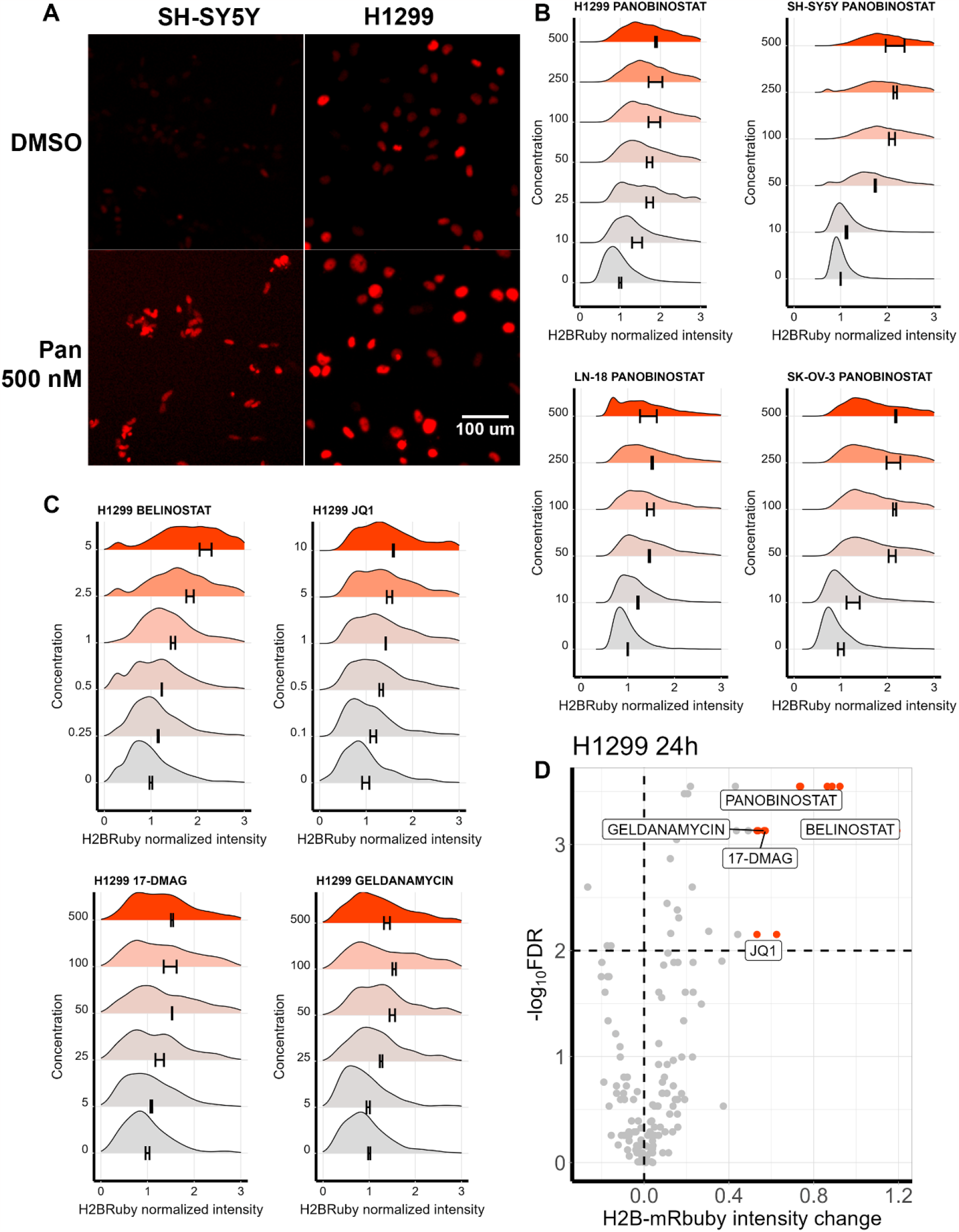
Chromatin remodulating drugs induce H2B-mRuby fluorescence. **A**. Representative images of nuclei H2B-mRuby fluorescence in H1299 and SH-SY5Y cells treated with DMSO or 500 nM panobinostat (Pan) for 24h. **B**. Distribution of median H2B-mRuby fluorescence per nucleus for H1299, SH-SY5Y, LN-18 and SK-OV-3 cells treated with panobinostat for 24h. H2B-mRuby fluorescence was normalized to cells treated with DMSO (zero concentration) and distribution was calculated based on average on 550 nuclei. On average three repeats were performed for each condition and 4 automatically selected fields were imaged. SD is indicated as +-range from mean based on mean fluorescence values for each repeat. **C**. Distribution of median H2B-mRuby fluorescence per nucleus for H1299cells treated with belinostat, JQ-1, 17-DMAG and geldanamycin for 24h. **D**. Volcano plot showing differential increase in H2B-mRuby fluorescence for H1299 cells treated with 30 different drugs for 24h compared to DMSO-treated cells. Each dot represents mean fluorescence for a drug used in particular concentration, maximum effects for statistically significant drugs are labeled. P-values were calculated using Mann-Whitney test based on mean values for each image and then Benjamini-Hochberg correction for multiple testing was applied (FDR).

## Discussion

Our results show, that although XTT assay and imaging methods often produce different results, the use of AUC metric overall provides consistent comparisons. Although IC50 is a convenient metric which offers an easily interpretable value it should be used with caution when comparing values from different datasets, especially obtained by different methods. We propose that it is more reliable to compare either concentrations which reduce proliferation by more than 20% or use AUC metric. The limitations of IC50 calculation can be somewhat bypassed by considerable increasing the number of drug concentrations used in a test, however this can significantly increase the cost and time for large-scale tests. Our findings are consistent with the results of other studies, which show that AUC or other area-based metrics, like DSS, produce more reliable results, especially for prediction of drug sensitivity^34,35^.

The difference between measurements performed by different methods also depended on a drug selection, especially for drugs that cause cell cycle arrest or other cytostatic drugs as has been shown before^18,36^. For example, palbociclib is known to induce cell size growth^37^ and accumulation of mitochondria, thus resulting in false results obtained by methods relying on mitochondrial activity^38^. We showed that palbociclib as well as other cell cycle inhibitor not only induce mitochondria accumulation due to increased cell size, but also increase mitochondrial activity itself. This effect we observed not only for well-established cell cycle inhibitors such as palbociclib and alisertib, but also for PARP inhibitor talazoparib and VEGFR inhibitor axitinib. Both inhibitors can induce DNA damage response and lead to G2/M arrest in some cell types^39,40^, which leads to senescence^41,42^ and probably to increased mitochondrial activity. High-content imaging allows to accurately distinguish effects on cell numbers, mitochondria accumulation and changes in mitochondrial activity. For example, dasatinib reduces cell size, which results in slightly decreased overall mitochondria signal, however dasatinib-treated cells had higher mitochonrdrial activity per occupied area, suggesting potentially different mechanisms of dasatinib action compared to similar inhibitors such as bosutinib^43^.

One of the concerns with the use of H2B-mRuby to count cells is that cells have heterogeneous levels of transgene expression, and since not all cells have detectable transgene expression the changes in the number of H2B-mRuby positive cells might not represent the changes in numbers of all cells. Also, the introduction of transgene might make transduced cell subpopulation more or less sensitive to a specific drug. However, even though our cell lines had varied percentage of H2B-mRuby positive cells (from 30 to 80%) and H2B-mRuby intensity, the results between nuclei counting with H2B-mRuby were highly consistent with nuclei counting using Hoechst staining or with results of trypan blue exclusion assay. The main drawback of using H2B-mRuby is the necessity of creating transgene cells, which might not be possible in case of *ex vivo* drug screens using patient-derived cells.

We describe several advantages of using H2B-mRuby: the ability to non-invasively record cell proliferation dynamics and find potential chromatin modulators. The proliferation dynamics can improve drug classification based on whether they prevent cell proliferation completely, reduce the initial cell numbers or allow cells to proliferate even at slower rates. We showed that H2B-mRuby intensity changes in response to chromatin remodeling drugs, such as HDAC and BET inhibitors. Several studies also suggested the possibility to use transgene expression to detect drugs that affect cell epigenetics. These approaches used cells with silenced GFP transgene and then drugs which affected epigenetic factors reactivated GFP expression, increasing the number of GFP positive cells^44-46^. We show that similar approach works even if introduced transgene was not completely repressed in the majority of cells, and chromatin remodelers can be identified by increase in overall transgene fluorescence. In our screen we detected increase in H2B-mRuby levels after treatment with HSP90 inhibitors. Although HSP90 inhibitors are not considered as classical chromatin remodelers there are reports that HSP90 can directly affect chromatin state^47^ and that HSP90 inhibitors such as 17-DMAG can also inhibit histone lysine demethylases^48^.

In conclusion, modern imaging-based approaches provide several benefits to large-scale drug screens, such as higher cell counting accuracy, ability to measure cell proliferation dynamics, and perform additional measurement such as the use of H2B-mRuby fluorescence intensity as reporter for chromatin remodulation. We show that AUC metric provides more consistency when comparing cell viability results obtained by imaging-methods with results of conventional assays. We also identify main reasons for measurement differences, such as increased cell size, induction of senescent phenotype or altered mitochondrial activity-factors, which should be considered for consistent integration of imaging data with existing large-scale drug screens.

## Methods

### Cell culture, lentiviral transduction and materials

All cell lines are not in the list of commonly misidentified cell lines that are controlled by the International Cell Line Authentication Committee. Table S1 contains list of reagents used for cell cultivation, cell densities used in experiments and growth conditions for each cell line. Cells were routinely checked for mycoplasma with Hoechst-33342 and DAPI staining, and to prevent mycoplasma contamination cells were treated with EZkillTM Mycoplasma Elimination Kit (HiMediaLabs) after defreezing. Lentiviral preparation was performed as described previously using pLentiPGK Hygro DEST H2B-mRuby2 (Addgene #90236)^49^ and then cells were transduced to achieve at least 30% transduction rate. Cells expressing H2B-mRuby were then enriched using selection with 0.5 mg/ml hygromycin b (Sigma) and verified using fluorescence microscopy and flow cytometry (LSRFortessa flow cytometer, BD Biosciences). All materials used, their manufacturers are listed in Table S1.

### Fluorescence microscopy

Cells were imaged on 96-well plates using motorized Leica DMI8 fluorescence microscope (Leica, Germany). For nuclei staining 1 μg/ml Hoechst-33342 was added, for tubulin and mitochondria imaging 1 μl of Tubulin Tracker™ Deep Red (Invitrogen, USA) and 100 ng/ml TMRE (Lumiprobe, Russia) were added, and then cells were incubated at 37°C and 5% CO_2_ for 30 minutes before imaging. For each well (biological repeat of drug treatment) six fields were automatically selected with the same pattern for all wells. For nuclei staining autofocus was performed on Hoechst-33342 images and for H2B-mRuby protein using bright field images. Plate layout and autofocus were done using LAS X software. Nuclei segmentation, illumination correction and measuring H2B-mRuby intensity were performed in CellProfiler v4.2. Cell segmentation for different cell sizes using nuclei, tubulin and mitochondria staining were performed using combination of Cellpose and Cellprofiler, and detailed protocol is available upon request. Image quality control was performed manually by two researches and using custom Python scripts. Images with low quality or presence of optical obstacles (areas with high background signal and serum debris) were excluded from analysis.

### Analysis of cell survival, IC50 and AUC calculation

Trypan blue exclusion assay was performed manually by two independent researchers using Neubauer chamber. Prior cell counting in Neubauer chamber cells were washed with PBS, trypsinized at 37°C and 5%CO_2_ for 5 minutes and resuspended in complete medium. XTT assay was measured by 450 nm absorbance and 650 nm reference using Multiskan FC (ThermoScientific, USA) after 4h incubation at 37°C and 5% CO_2_, reference signal for each well and mean signal for wells containing only growth medium and XTT were subtracted before normalization. Number of nuclei for each image was calculated using CellProfiler pipeline and data was extracted using custom Python script. All experiments were performed in at least three replicates. Details for drug concentrations, manufacturers and storage conditions is provided in Table S1. In cases when drugs were dissolved in DMSO, the amount of DMSO was made equal for all drug concentrations including control treatment (no drug added). For all experiments the amount of DMSO was kept below 0.1%. All results were normalized to mean values of control treatment (no drug), control treatment was considered as 100% and complete absence of cells as 0%. For cell dynamics number of nuclei were also normalized to the number of nuclei at the start of experiment for each repeat. IC50 values were calculated using four-variable non-linear regression in GraphPad Prism 9 software with top and bottom values set at 100 and 0. AUC values were calculated using trapezoidal rule in Python. AUC values for large-scale drug screens for GDSC1/2^3^, CTRP^1^ and PRISM^6^ datasets was downloaded from DepMap^50,51^ website (https://depmap.org/portal/).

## Statistical analysis

Mann-Whitney test and t-tests were performed using SciPy Python package and Benjamini-Hochberg correction for multiple testing was performed using statmodels Python package. Spearman’s correlation was calculated using GraphPad Prism 9 and SciPy Python package. Mean, SEM and SD values for cell viabilities were calculated in R.

## Data availability

All data, codes and pipelines are available upon request and will be made publicly available upon publication.

## Acknowledgements

Cell imaging was supported by RSF grant 23-74-10103. Identification of chromatin modulating drugs using H2B-mRuby fluorescence was supported by RSF grant 22-14-00353.

## Author Contributions

Mikheeva A.M.: conceptualization, original draft preparation, investigation, data curation, formal analysis, methodology, resources, visualization; Bogomolov M.A.: investigation, methodology, software, validation; Sementsov M.V.: investigation, validation; Spirin P.V.: review and editing, funding acquisition; Prassolov V.S.: review and editing, funding acquisition, supervision; Lebedev T.D.: conceptualization, original draft preparation, investigation, data curation, formal analysis, methodology, visualization, software, project administration, funding acquisition, and supervision.

